# A functional triangle relationship of topoisomerase IIβ, BARD1, and ERK2 in immediate early gene transcription

**DOI:** 10.64898/2026.04.20.719704

**Authors:** Heeyoun Bunch, Jaehyeon Jeong, Anh Cong, Jaeyeon Jang, Inuk Jung, Keunsoo Kang, Matthew J. Schellenberg

**Affiliations:** Department of Applied Biosciences, Kyungpook National University, Daegu 41566, Republic of Korea; School of Applied Biosciences, College of Agriculture & Life Sciences, Kyungpook National University, Daegu 41566, Republic of Korea; Department of Biochemistry and Molecular Biology, Mayo Clinic, Rochester, MN 55905, USA; School of Computer Science and Engineering, Kyungpook National University, Daegu 41566, Republic of Korea; Department of Microbiology, College of Bio-convergence, Dankook University, Cheonan 31116, Republic of Korea

**Keywords:** Transcriptional regulation, topoisomerase IIβ, BARD1, ERK2, immediate early genes, RNA polymerase II transcription, gene regulation

## Abstract

Immediate early genes (IEGs) encode master transcription factors, including *EGR1*, *FOS*, *JUN*, and *MYC* genes that drive robust transcription in the G1 phase of cell cycle. Our previous studies indicate that topoisomerase IIβ (TOP2B) critically regulates IEG transcription and that the activities of TOP2B are dynamically controlled through post-translational modifications by BRCA1-BARD1 and ERKs.

Here, we show that two E2 enzymes, UBCH5b and UBC13/MMS2, differentiate the effects and functions of TOP2B ubiquitination by BRCA1-BARD1. Comprehensive transcriptomics, proteomics, and biochemical and molecular cellular analyses revealed a close relationship between BARD1 and key mitogen-activated protein kinase pathway genes and identified activated ERK2 as a novel kinase that phosphorylates BARD1 at S391, a previously reported mitotic phosphorylation site, whose genetic mutation has been linked to tumorigenesis. Mechanistically, the catalytic activity of ERK2 stimulates TOP2B ubiquitination mediated by BRCA1-BARD1 in complex with UBCH5b and UBC13/MMS2, which controls the binding and function of TOP2B and BARD1 for transcriptional activation at representative IEGs. Taken together, our data propose that there is a functional regulatory circuit involving TOP2B, BARD1, and ERK2, three key transcriptional activators for IEG transcription, in which the gene association and catalytic activity of TOP2B are regulated through E2-differentiated ubiquitination by BRCA1-BARD1 and the phosphorylation of BARD1 by ERK2 for productive transcription.

## BACKGROUND

Metazoan transcription of messenger and most non-coding RNAs (mRNAs and ncRNAs) is carried out by a sole RNA polymerizing enzyme, RNA polymerase II (Pol II), and numerous transcription factors that act across the stages of transcriptional initiation, early elongation featuring promoter-proximal pausing/pause release, productive elongation, and termination[1–4]. These transcription factors modulate Pol II association/dissociation, polymerization and post-disengagement RNA cleavage, pausing and forward-movement, and catalytic rate[5–8]. Transcription factors and Pol II form complex, interactive and dynamically evolving networks, enabling reciprocal gene regulation in response to internal and external stimuli and cellular needs. Accumulating evidence suggests that categorizing transcriptional activators or repressors is context-dependent rather than absolute, as many transcriptional factors have biphasic functions, even within a single gene, that depend on protein modifications or binding partners[9–12]. Post-translational modifications (PTMs), covalently and reversibly attaching functional inorganic or organic compounds to protein factors, are considered major molecular switches to divert functions. More than 200 distinctive PTMs have been reported, with phosphorylation and ubiquitination among the most prominent ones[13].

Protein phosphorylation is mediated by protein kinases that transfer γ-phosphates from ATP to their substrate proteins. Phosphorylation often functions as a molecular switch to turn on and off enzymatic activities, regulates inter- and intra-molecular interactions, and determines subcellular localization and protein stability[14–16]. Ubiquitin is a small regulatory peptide (8.6 KDa), abundant and ubiquitously present in eukaryotes. Ubiquitination of a substrate protein is mediated by sequential reactions of three enzymes: an E1 ubiquitin-activating enzyme, an E2 ubiquitin-conjugating enzyme, and an E3 ubiquitin ligase[17]. There are eight E1 enzymes, approximately 40 E2 enzymes, and over 600 E3 enzymes in humans[18]. Ubiquitination is an established signal for proteasome-mediated proteolysis, in which K48-linked poly-ubiquitination tags are attached to destinate misfolded, aged, or targeted proteins for degradation[19]. Short-chained ubiquitination or K63-linked polyubiquitination has been reported to regulate transcription, DNA repair, molecular interaction, translocation, and protein activity[19, 20]. Recently, one study indicated the presence of non-canonical ubiquitination by E2s, conjugating ubiquitin to serine/threonine, instead of lysine of target proteins, without requiring E1 and E3 enzymes[21]. Together with phosphorylation, protein ubiquitination is a crucial molecular event to maintain and perform cellular processes.

BRCA1 and BARD1 proteins form a heterodimeric complex that functions as a key DNA repair factor in homologous recombination and non-canonical non-homologous end joining as well as an E3 ubiquitin ligase[22–25]. Double-stranded DNA breakage (DSB) induces DNA damage response (DDR) signaling, which activates the PI3K-like kinases, ataxia telangiectasia mutated (ATM), DNA-dependent protein kinase (DNA-PK), and ataxia telangiectasia and Rad3-related (ATR)[26–28]. These kinases phosphorylate various DNA repair, cell cycle, and transcription factors, including BRCA1, TRIM28, p53, Chk1/2, γH2AX, and WRN[10, 26, 27, 29–32]. BRCA1-BARD1 facilitates DNA resection of the broken DNA end for homologous-recombination[23]. Known ubiquitination substrates of the BRCA1-BARD1 E3-ligase activity include H2A, PCNA, NF2, estrogen receptor α, and TOP2B[22, 29, 33–35]. Importantly, BRCA1-BARD1 is a transcriptional activator of a subset of genes including NF-κB-responsive genes and IEGs[29, 36–39]. Our previous study showed that BRCA1-BARD1 ubiquitinates TOP2B and that BARD1 interacts with TOP2B with high affinity[29]. Additionally, either BRCA1 or BARD1 knockdown (KD) destabilizes TOP2B[29]. Ubiquitinated TOP2B has a stronger affinity for the DNA fragments encompassing the *EGR1* transcription start site (TSS) in comparison with ubiquitin-stripped TOP2B, which suggests the role of BRCA1-BARD1 in the regulation of TOP2B genome association[1, 29]. Supporting this finding, BRCA1 and TOP2B are co-localized genome-wide, suggesting that they collaborate for gene regulation[1, 29].

For IEG transcriptional activation, proliferating signals are transmitted from ligand-bound cell-surface receptors to nuclear target genes through the canonical mitogen-activated protein kinase (MAPK) signaling pathway[40]. A series of phosphorylation-based kinase activation cascades promote the phosphorylation/activation of ERK1 (MAPK3) and ERK2 (MAPK1) and their translocation from the cytoplasm to the nucleus[40]. Then, activated ERK2 phosphorylates and activates ELK1, a potent DNA binding transcription factor, to stimulate transcription of its target genes (e.g. *EGR1*)[14, 40].

Notably, IEGs are representative examples of stress-inducible genes, where Pol II is paused in the TSSs before the reception of the activating signal[30, 41–43]. Interestingly, transcriptional burst and Pol II pause release in IEGs are accompanied by and require TOP2B-mediated DDR signaling, featuring the phosphorylation and activation of ATM, DNA-PK/KU70/KU80, ATR, TRIM28, γH2AX, and BRCA1-BARD1[29, 30, 43, 44]. It is still unclear what triggers TOP2B-mediated DNA breakage and yet, the cryo-EM structure of the etoposide-poisoned TOP2B-*EGR1* TSS complex has revealed dramatic DNA bending around a DSB, altering local DNA topology and presumably relieving DNA torsion or regulating DNA winding/helicity[14]. Consistent with this conjecture, transcriptional activation alters DNA supercoiling profiles, and DNA positive or negative supercoiling reportedly activates or represses transcription[11, 45–51]. Furthermore, TOP2B knockout (KO) renders the genome to be more negatively supercoiled, underwound, and less accessible, thus deregulating transcription-mediated genomic topological changes[1, 11].

Recent studies, including ours and those of others have revealed distinct functions exerted by ERK1 and ERK2 despite about 85% identical protein primary sequences with less conserved N- and C-terminal domains (NTD and CTD)[14, 52–55]. Biochemical analyses demonstrated that both ERK1 and ERK2 phosphorylate the CTD of TOP2B targeting both shared and unique residues[14]. ERK1 stimulates TOP2B catalysis of both positive and negative supercoiling[14]. Interestingly, ERK2 retards TOP2B catalysis of transcriptionally favorable negative supercoiling, whereas it enhances TOP2B resolution of positive supercoiling[14]. In representative IEGs, ERK2 functions as a transcriptional activator, while ERK1 acts as a transcriptional repressor[14]. These findings are well-aligned with the reportedly distinctive roles of ERK1 and ERK2, in which only ERK2 is pro-proliferative[52–54].

In this study, we screened a panel of 11 human E2 enzymes and identified E2 enzymes, UBCH5b and UBC13/MMS2 that function with BRCA1-BARD1 E3 ligase to ubiquitinates TOP2B, yet in substantially distinct fashions. Interestingly, depleting UBCH5b or UBC13 also displays differential effects on TOP2B. UBCH5b KD increases cellular TOP2B level, while reducing genome-associated TOP2B, which suggests a role in regulating TOP2B-genome interaction and TOP2B destabilization. UBC13 KD markedly decreases TOP2B catalytic activities, as well as both cellular and chromosome-bound TOP2B protein levels. KD of either E2 enzymes or BARD1 represses IEG transcription. We found that BARD1 determines protein stability of TOP2B, UBCH5b, and UBC13 and the transcriptional output of a number of genes. Importantly, deletion of a single BARD1 allele paradoxically increases key factors in MAPK pathway, including *RAF1*, *JNK*s, and *ERK*s.

Mechanistically, our biochemical and mutational analyses revealed that BARD1 is phosphorylated at residue S391 by constitutively active ERK2 (ERK2m), and that ERK2 KD or catalytic inhibition results in abnormal increases of TOP2B and BARD1 occupancies and transcriptional suppression at representative IEGs. ERK2m enhances TOP2B ubiquitination by BRCA1-BARD1 with UBCH5 or UBC13/MMS2, suggesting that catalytically-active ERK2 phosphorylates BARD1 at S391 and promotes BRCA1-BARD1-mediated TOP2B ubiquitination to regulate TOP2B catalysis (UBC13/MMS2) and association with the DNA (UBCH5b) for transcriptional activation in IEGs.

## METHODS

### Cell culture and conditions

Mammalian cell culture and serum starvation and induction with HEK293 cells were carried out as previously described. An ERK2 inhibitor, VX-11e (Selleck, S7709) was used at a final concentration of 100 nM in 0.1% DMSO for 3 h. Control cells were treated with the same amount of DMSO for the same duration. CRISPR-Cas9 genome editing was performed to generate a BARD1 KO HEK293 cell line (Ubigene Biosciences). Homozygous KO (BARD1 deletion in both alleles) HEK293 cells were unviable, as previously reported[56], and a heterozygous (BARD1 deletion in a single allele) KO cell line was achieved and used in this study. Briefly, for BARD1 KO, the gRNAs were designed using the online CRISPR design tool (Red Cotton, https://en.rc-crispr.com/). The exon 4 region of BARD1 was selected to be targeted by CRISPR/Cas9 genome editing. A ranked list of gRNAs was generated with specificity and efficiency scores. The pair of oligos for two targeting sites was annealed and ligated to the YKO-RP006 vector (Ubigene Biosciences).

The YKO-RP006-hRABL6[gRNA] plasmids containing each target gRNA sequences were transfected into cells with Lipofectamine 3000 (Thermo Fisher Scientific). One or two days after the transfection, puromycin were added to screen the cells. After antibiotic selection a certain number of cells were diluted by limited dilution method and inoculated into 96-well plate. Selection of single clones were performed after 2–4 weeks and selected BARD1 KO clones were validated by PCR and Sanger sequencing. The gRNAs and primers for CRISPR design are shown in Supplementary Table (**Table S1**).

### Cloning and protein preparation

A human BRCA1 expressing vector, pcBRCA1-385 was originally gifted from Dr. Mike Erdos at the National Institutes of Health, and was cloned into pET17b in our previous study[29]. A full length BARD1 cDNA cloned into pCMV3-C-His was obtained from Sino Biological (HG15850-CH) and the BARD1 coding sequence was cloned into pET29b in our previous study[29]. For protein over-expression in BARD1 heterozygous knock-out HEK293 cells, BARD was conjugated with green fluorescent protein (GFP), generating pCMV3-His-GFP-BARD1. S391A and S391D mutations of BARD1 in pCMV3-His-GFP-BARD1 were introduced using specific primer sets, following the principle of Quikchange mutagenesis. A GFP expressing pCMV3 was cloned as a control. Constructed plasmids and cloning products were validated by sequencing. All primers used in this study, unless stated otherwise, were purchased from Integrated DNA Technology (**Table S1**).

Human ERK1/R84S ERK1 (ERK1m, constitutively active mutant) and ERK2/R67S ERK2 (ERK2m, constitutively active mutant) were bacterially expressed and purified as described in our previous work[14]. Ubiquitin- and phosphate-stripped TOP2B was purified from HEK cells and validated by an *in vitro* decatenation assay and immunoblotting, as previously described[11, 14, 29]. Full-length BRCA1 and BARD1 constructs were PCR amplified from bacterial expression vectors, pET-WT-BRCA1[29] and pET29b-WT-BARD1[29] using primer sets (**Table S1**) and cloned into pMCentr2 vector [DNASU] using Ligation Independent Cloning[57]. Each protein construct in pMCentr2 vector was recombined into a destination mammalian protein expressing vector using Gateway Cloning.

Specifically, BARD1 was recombined into pcDNA6.2/N-YFP-Dest (Thermofisher) and BRCA1 was recombined into N-mCherry-Dest. All cloning steps and plasmid propagation were carried out in XL10-Gold Ultracompetent Cells (Agilent) and plasmid inserts were verified by sequencing. A giga-prep kit (Zymo Reserch) was used to prepare milligram-scale DNA, and plasmids were further purified using phenol-chloroform extraction and ethanol precipitation prior to being redissolved in nanopure water for transfection. Purified plasmids encoding BRCA1 and BARD1 were used to transfect HEK293F cells using a protocol described previously[58, 59]. Briefly, a 1.2 mg mixture of BRCA1 and BARD1 plasmids (3:2 ratio) was combined with 1.4 mg PEI (Polysci) and transfected into 1.2x10^9^ HEK293F cells in 500 mL of Hycell TransFx-H media (Cytiva). Cells were grown for 72 h and fluorescence was measured every 24 h post transfection to monitor protein expression. Cells were harvested by centrifugation at 400 g and 4 °C for 10 min, then resuspended in 50 mL PBS before being pelleted again at 500 g for 10 min. Cells were lysed in 36 mL of cell lysis buffer containing 50 mM Tris pH 8.0, 600 mM NaCl, 0.5% (v/v) NP-40 substitute (Sigma) and 1 mM TCEP, supplemented with Complete-EDTA free protease inhibitor cocktail (Roche). Lysate was sonicated for 3 cycles of 15 s sonication at 50% power using a Branson SFX250 sonicator, with a 30-second rest period between each cycle. Clarified lysate was collected by centrifugation at 25,000 g and 4 °C for 15 min, then passed over a column with a 2 mL bed volume of pre-equilibrated LaM4 anti-mCherry sepharose resin. Protein-bound resin was washed twice with lysis buffer and three times with wash buffer (50 mM Tris pH 8.0, 400 mM NaCl, 0.1% NP-40, 0.5 mM TCEP, and 0.1x Complete-EDTA free protease inhibitor cocktail). The resin was subjected to three rounds of low salt wash using buffer containing 20 mM Tris pH 8.0, 300 mM NaCl, and 1 mM TCEP, following by a final 2.5 mL of low salt wash containing 20 μg/mL TEV protease, then stoppered for incubation overnight at 4 °C. BRCA1-BARD1 complex was eluted from the resin using 6 mL of size-exclusion chromatography buffer containing 20 mM Tris pH 7.5, 500 mM NaCl, and 0.5 mM TCEP. The elution was concentrated to 2 mL volume using an Amicon 10K cutoff centrifugal filter (Millipore) then loaded onto Superdex 200 Increase 10/300 size-exclusion chromatography column (Cytiva). Fractions containing BRCA1-BARD1 complex and BARD1 proteins were separated in the fractions and visualized on silver-stained SDS-PAGE gels. The final proteins were concentrated and stored in a buffer solution containing 20 mM Tris pH 7.5, 300 mM NaCl, 1 mM TCEP, 25 % (v/v) glycerol and 0.01x complete protease inhibitor.

### Real-time PCR

Total RNA was purified using the RNeasy kit (Qiagen) following the manufacturer’s instructions. The concentration of RNA was measured using Nanodrop. cDNA construction and real-time quantitative PCR was conducted, as previously described and with the indicated primers (**Table S1**). The results were presented as relative fold differences, standard deviations (SDs), and statistical validations (see below).

### RNA-seq and bioinformatics analysis

WT and BARD1 heterozygous KO HEK293 cells were grown to approximately 70% confluence and collected by scrapping. Total RNAs were extracted from the cells and purified using RNeasy kit (Qiagen). cDNA construction and sequencing were performed by Rokit Genomics (South Korea). The libraries were prepared for 151PE sequencing using TruSeq stranded Total RNA Kit (Illumina). Ribosoamal RNA is removed using biotinylated, target-specific oligos combined with Ribo-Zero rRNA removal beads from 1 μg of total RNA. Rest of RNA molecules were purified and fragmented. The fragmented RNAs were synthesized as single-stranded cDNAs through random hexamer priming. By applying this as a template for second strand synthesis, double-stranded cDNA was prepared. After sequential process of end repair, A-tailing and adapter ligation, cDNA libraries were amplified with PCR. Quality of these libraries was estimated with TapeStation 4200 instrument and D1000 ScreenTape System (Agilent). They were quantified with the KAPA library quantification kit (Kapa Biosystems) according to the manufacturer’s library quantification protocol. Following cluster amplification of denatured templates, sequencing was progressed as paired-end (2 × 151bp) using Illumina NovaSeq X plus (Illumina). We first processed raw paired-end RNA-seq reads with fastp (v0.23.4)[60] to trim adapter sequences and remove low-quality bases. For paired-end libraries, default adapter-detection settings were used, and reads with a Phred quality score below 20 or a length shorter than 30 bp were discarded. The filtered reads were then aligned to the human reference genome (GRCh38) using STAR (v2.7.11b)[61], and coordinate-sorted BAM files were generated for downstream analyses. Gene-level read counts were obtained with featureCounts (v2.0.8)[62] using the GENCODE human release 44 annotation for GRCh38.p14 (https://www.gencodegenes.org/human/releases.html)[63]. Read counting was performed in paired-end mode with the -p and --countReadPairs options. Gene IDs were kept as the primary feature identifiers, while gene_name and gene_type were retained as additional annotations. We performed differential expression analysis using PyDESeq2 (v0.5.4)[64], a Python implementation of the DESeq2 framework[65]. Genes with fewer than 10 counts in more than four samples were removed before analysis. The filtered count matrix was analyzed with the design formula ∼ condition, and differential expression between BARD1^−/+^ and WT HEK293 cells was assessed using the Wald test. *P*-values were adjusted for multiple testing with the Benjamini-Hochberg method[66]. Genes with an adjusted *P*-value < 0.05 and an absolute *Log2* fold change > 1 were considered differentially expressed. For exploratory analyses, we used normalized expression values for principal component analysis (PCA) and sample correlation analysis. PCA was performed after size-factor normalization and *Log2* transformation using the most variable genes, and sample-to-sample similarity was evaluated by Pearson correlation. For heatmap visualization of differentially expressed genes, expression values were *Log*_2_-transformed and Z-score normalized across samples. Heatmaps were generated using pheatmap (v1.0.13) with hierarchical clustering based on Euclidean distance and complete linkage. KEGG pathway enrichment analysis was performed using clusterProfiler (v4.18.4) with an adjusted *P*-value cutoff of 0.05. The Homo sapiens KEGG reference database was used for pathway annotation.

### Immunoblotting

The procedure and reagents for immunoblotting were identical to those used in our previous studies[10, 14, 29, 30]. The primary antibodies used in this study were BARD1 (sc74559, Santa Cruz Biotechnology; A300-263A, Bethyl Laboratories), BRCA1 (A300-000A, Bethyl Laboratories), UBCH5B (LS-C352970-100, LSBio), UBC13 (#4919S, Cell Signaling Technology), ERK2 (ab32081, Abcam), γH2AX (sc517348, Santa Cruz Biotechnology), TOP2A (sc365916, Santa Cruz Biotechnology), α-Tubulin (sc8035, Santa Cruz Biotechnology), ACTIN (MA5-15739, Invitrogen), ubiquitin (sc8017, Santa Cruz Biotechnology), and TOP2B (A300-949A and A300-950, Bethyl Laboratories; sc-25330, Santa Cruz Biotechnology).

### Chromatin immunoprecipitation and qPCR

ChIP-qPCR analysis was conducted following the previously described methods and reagents without modifications[11, 14, 29, 30]. The antibodies used for this study were phosphorylated S2 Pol II (ab5095, Abcam), BARD1 (A300-263A, Bethyl Laboratories), TOP2A (sc365916, Santa Cruz Biotechnology), and TOP2B (A300-949A and A300-950, Bethyl Laboratories). The primer sets used are listed in **Table S1**.

### *In vitro* kinase assay

The substrate, 1 μg of BARD1 was incubated with approximately 0.2–0.5 μg WT or mutant ERK proteins as a kinase in a reaction. For the control reaction, BARD1 without any kinase (ERK storage buffer only) was prepared side-by-side. Kinase buffer included 25 mM Tris pH 8.0, 2 mM DTT, 500 μM cold ATP, 100 mM KCl, 10 mM MgCl_2_, and 2.5 μCi γ-P^32^ (PerkinElmer) labeled ATP to be final concentrations per a reaction[14]. The reaction was incubated at RT for 1 h before it was terminated using an 8× SDS-loading buffer. The reaction was subjected to SDS-PAGE followed by autoradiography. For mass spectrometry, cold ATP without γ-P^32^ labeled ATP was used and the reaction was allowed for 1 h at 30 °C and separated on a 7% SDS-PAGE gel. The bands corresponding to BARD1 were sliced for the further analysis.

### ICE assay

ICE assay was performed using the Human Topoisomerase ICE Assay Kit (TG1020-2b, Topogen), according to the manufacturer’s instruction. Cells were grown to about 60% confluence in 6-well plates, subjected to mock or siRNA-mediated KD treatment for 48–72 h, and supplied with DMSO or 50 μM etoposide for 1 h, prior to harvesting the cells. Genome-associated TOP2A and TOP2B were quantified by antibodies: A300-950A, Bethyl Laboratories for TOP2B and sc-365916, Santa Cruz Technologies for TOP2A.

### Post-translational modification analysis

Each gel slice was subjected to in-gel digestion with trypsin. The resulting peptides were dried and reconstituted in 0.1% trifluoroacetic acid (TFA) and 2% acetonitrile. For ubiquitination site analysis, peptides from control TOP2B and BRCA1/BARD1 samples were analyzed by LC–MS/MS without further enrichment. For TOP2B and BRCA1/BARD1 complexes treated with UBCH5b or UBCH13, K-ε-GG–modified peptides were enriched from tryptic digests using the PTMScan® HS Ubiquitin/SUMO Remnant Motif Kit (Cell Signaling Technology) according to the manufacturer’s instructions. The enriched peptides were desalted using in-house C18 StageTips, dried, and reconstituted in 0.1% TFA and 2% acetonitrile prior to LC–MS/MS analysis. Phosphorylation site analysis of BARD1 was performed using the High-Select Fe-NTA Phosphopeptide Enrichment Kit (Thermo Scientific) according to the manufacturer’s instructions. The enriched phosphopeptides were desalted using in-house C18 StageTips and analyzed by LC–MS/MS.

### LC–MS/MS analysis

Mass spectra were acquired using a Thermo Scientific LTQ-Orbitrap Velos Pro mass spectrometer coupled to a nano-flow UHPLC system (ADVANCE UHPLC; AMR Inc.) equipped with an Advanced Captive Spray source (AMR Inc.). Peptide mixtures were loaded onto a C18 trap column (PepMap Neo Trap Cartridge, 0.3 mm × 5 mm, 5 μm particle size; Thermo Fisher Scientific) and subsequently separated by C18 reverse-phase chromatography (0.075 mm × 150 mm, 3 μm particle size; CERI). Peptides were eluted with a linear gradient of solvent B (5–35% acetonitrile containing 0.1% formic acid) at a flow rate of 300 nL/min over 60 min. Solvent A consisted of 0.1% formic acid in water, and solvent B consisted of 0.1% formic acid in acetonitrile.

For ubiquitinated peptide analysis, the mass spectrometer was operated in data-dependent acquisition mode with eleven successive scans, including a full MS scan (m/z 350–1,800) acquired in the Orbitrap at a resolution of 60,000, followed by MS/MS scans of the top 12 most intense precursor ions using collision-induced dissociation (CID) in the ion trap. MS/MS spectra were acquired with a normalized collision energy of 35%, an isolation width of 2 m/z, and a dynamic exclusion time of 90 s. For phosphopeptide analysis, the mass spectrometer performed seven successive scans, consisting of a full MS scan (m/z 350–1,600) acquired in the Orbitrap at a resolution of 60,000, followed by data-dependent MS/MS scans of the top three most abundant precursor ions. The second to fourth scans were performed using CID in the ion trap, and the fifth to seventh scans were performed using higher-energy collisional dissociation (HCD) in the Orbitrap at a resolution of 7,500. MS/MS spectra were acquired with a normalized collision energy of 35% and a dynamic exclusion time of 90 s.

Raw data files were searched against the UniProt Homo sapiens proteome database (downloaded October 2020) and the cRAP contaminant protein database using the MASCOT search engine (version 2.6; Matrix Science) via Proteome Discoverer 2.5 (Thermo Fisher Scientific).

Carbamidomethylation of cysteine was set as a fixed modification, while oxidation of methionine and acetylation of protein N-termini were set as variable modifications. For phosphopeptide identification, phosphorylation of serine, threonine, and tyrosine residues was included as variable modifications.

For ubiquitinated peptide identification, GlyGly modification of lysine residues was included as a variable modification. The maximum number of missed cleavages was set to two.

### Statistics and Reproducibility

Standard deviation was calculated and used to generate error bars. One-sided Student’s t-test or one-way ANOVA with multiple comparisons used to determine statistical significance (*P* < 0.05). Graphs were generated using the Prism 8 software (GraphPad, Inc.).

Representative experimental images of gels and autographs were repeated at least twice independently with similar results.

### Data Availability

All data are available in the manuscript or as supplementary information. Source data are provided with this paper. The proteomics data generated in this study have been deposited in ProteomeXchange under accession codes PXD077421 (phosphopeptide analysis) and PXD077423 (ubiquitinated peptide analysis) and jPOST under accession codes JPST004487 (phosphopeptide analysis) and JPST004488 (ubiquitinated peptide analysis). The RNA-seq data generated in this study have been deposited in the Gene Expression Omnibus Database under accession code GSE328750.

## RESULTS

### BRCA1-BARD1 assembles with the E2 enzymes, UBCH5b or UBC13/MMS2, to ubiquitinate TOP2B

To identify E2 enzymes that collaborate with the BRCA1-BARD1 complex to ubiquitinate TOP2B, 11 human E2 enzymes were screened by the *in vitro* ubiquitination and immunoblotting assays[29]. The degrees of TOP2B ubiquitination were quantified, compared with the control TOP2B from the reaction lacking an E2 enzyme. The data indicated that the E2 enzymes UBCH5b and UBC13/MMS2, together with BRCA1-BARD1 E3 ligase, ubiquitinate TOP2B (**Fig. 1A** and **Supplementary Figure 1A**). Next, we sought to identify the residues in TOP2B that were ubiquitinated by BRCA1-BARD1 with UBCH5b or UBC13/MMS2. Additionally, pre-existing ubiquitinated sites of the control proteins, TOP2B, BRCA1, and BARD1 were identified by in-gel digestion and liquid chromatography tandem mass spectrometry (LC–MS/MS)(**Fig. 1B** and **Supplementary Data 1–3**). TOP2B was gel-sliced after *in vitro* ubiquitination assays containing either E2 enzyme and was subsequently analyzed by in-gel digestion, K-GG enrichment, and LC–MS/MS. (**Figs. 1C–E**). Mass spectrometry data revealed strikingly distinct ubiquitination patterns, in particular for TOP2B, depending on the interacting E2 enzymes (**Figs. 1D,E**, **Supplementary Figure 1B,** and **Supplementary Data 1**). UBCH5b mediated ubiquitination of 31 residues throughout the ATPase, DNA binding & catalytic, and C-terminal domains of TOP2B. In contrast, UBC13/MMS2 in association with BRCA1-BARD1 ubiquitinated 5 residues, including K510, K744, and K979, only in the DNA binding & catalytic domain (**Fig. 1E** and **Supplementary Figure 1A**). Additionally, the unique and common 60 and 34 ubiquitinated residues in BRCA1 and BARD1 were identified and mapped on the proteins (**Fig. 1F**, **Supplementary Figure 1C,** and **Supplementary Data 2, 3**). These data suggested that both UBCH5b and UBC13/MMS2 induce BRCA1-BARD1 to be auto-ubiquitinated at mutual and distinctive sites and that UBCH5b dictates BRCA1-BARD1 to ubiquitinate its substrate TOP2B across a broader number of sites, while UBC13/MMS2 mediates BRCA1-BARD1 to ubiquitinate TOP2B at fewer, more specific sites in the DNA binding & catalytic domain.

**Fig. 1.**
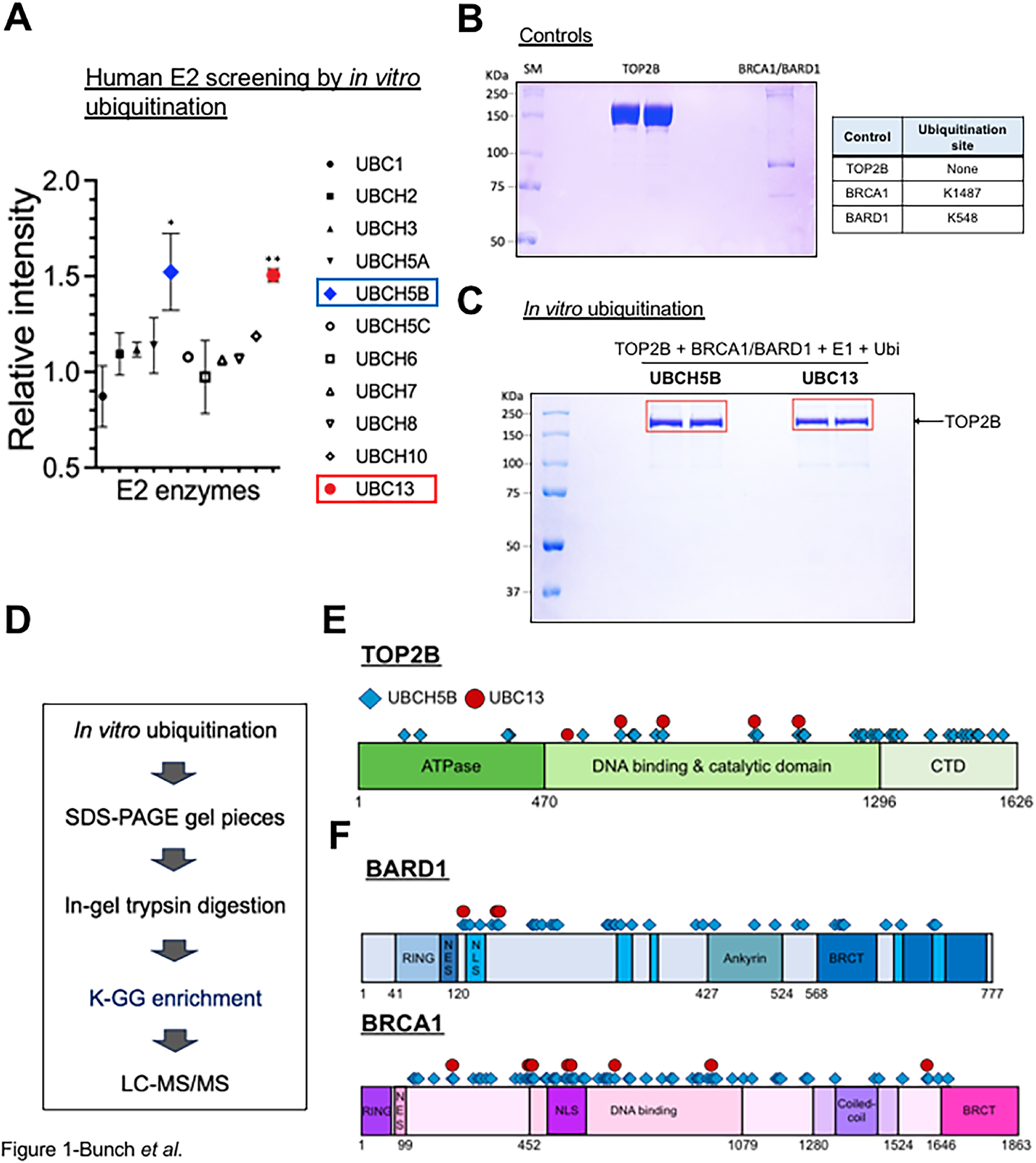
BRCA1-BARD1 assembles with E2 enzymes, UBCH5b or UBC13/MMS2, to ubiquitinate TOP2B. (**A**) Statistical summary of the *in vitro* ubiquitination assay and immunoblotting-based quantification results. A total of commercially available, 11 human E2 enzymes were screened, among which UBCH5b and UCB13/MMS2 were identified to interact with BRCA1-BARD1 (E3 ligase) and UBE1 (E1) to ubiquitinate TOP2B (substrate). (**B**) Coomassie-blue stained control TOP2B and BRCA1-BARD1 proteins used for the *in vitro* ubiquitination and LC-MS/MS analysis. Ubiquitinated residues identified in the control samples were listed next the gel image. SM, standard protein size marker. (**C**) Post-*in vitro* ubiquitinated TOP2B samples on a SDS-PAGE gel, which were sliced and analyzed by LC-MS/MS analysis. (**D**) A flow chart showing the major steps of mass spectrometry analyses to identify ubiquitinated sites of TOP2B by BRCA1-BARD1 in complex with either UBCH5b and UBC13/MMS2. (**E**) The residues identified to be ubiquitinated by UBCH5b-BRCA1-BARD1 (light blue diamonds, n = 35) or UBC13/MMS2-BRCA1-BARD1 (red circles, n = 5) were marked on TOP2B protein, showing distinctive patterns and specificities of TOP2B ubiquitination by BRCA1-BARD1, depending on partnering E2 enzymes. Note that the resides ubiquitinated by UBC13/MMS2-BRCA1-BARD1 were mapped only in the DNA binding and catalysis domain. Four sites, K643, K744, K979, and K1079 were mutually identified between two groups, with UBC13/MMS2-unique site, K510. (**F**) The residues identified to be ubiquitinated by UBCH5b-BRCA1-BARD1 (light blue diamonds) or UBC13/MMS2-BRCA1-BARD1 were marked on BARD1 and BRCA1 proteins (red circles).

### BARD1, UBCH5b, or UBC13 knockdown deregulates TOP2B and represses IEG transcription

To understand the function of TOP2B ubiquitination by the two E2-E3 entities UBCH5b-BRCA1-BARD1 and UBC13/MMS2-BRCA1-BARD1, each of BARD1, UBCH5b, and UBC13 was depleted from HEK293 cells using small interfering RNA (siRNA) species and the control RNA (scrambled RNA control, SCR) and the gene KD was validated by immunoblotting (**Fig. 2A**). Notably, we previously showed that BARD1 is important for TOP2B stability, as BARD1 KD decreases TOP2B protein levels[29]. Interestingly, UBCH5b KD significantly increased the cellular TOP2B protein level, whereas UBC13 KD mildly yet consistently decreased it (**Fig. 2B** and **Supplementary Figure 2A**).

**Fig. 2.**
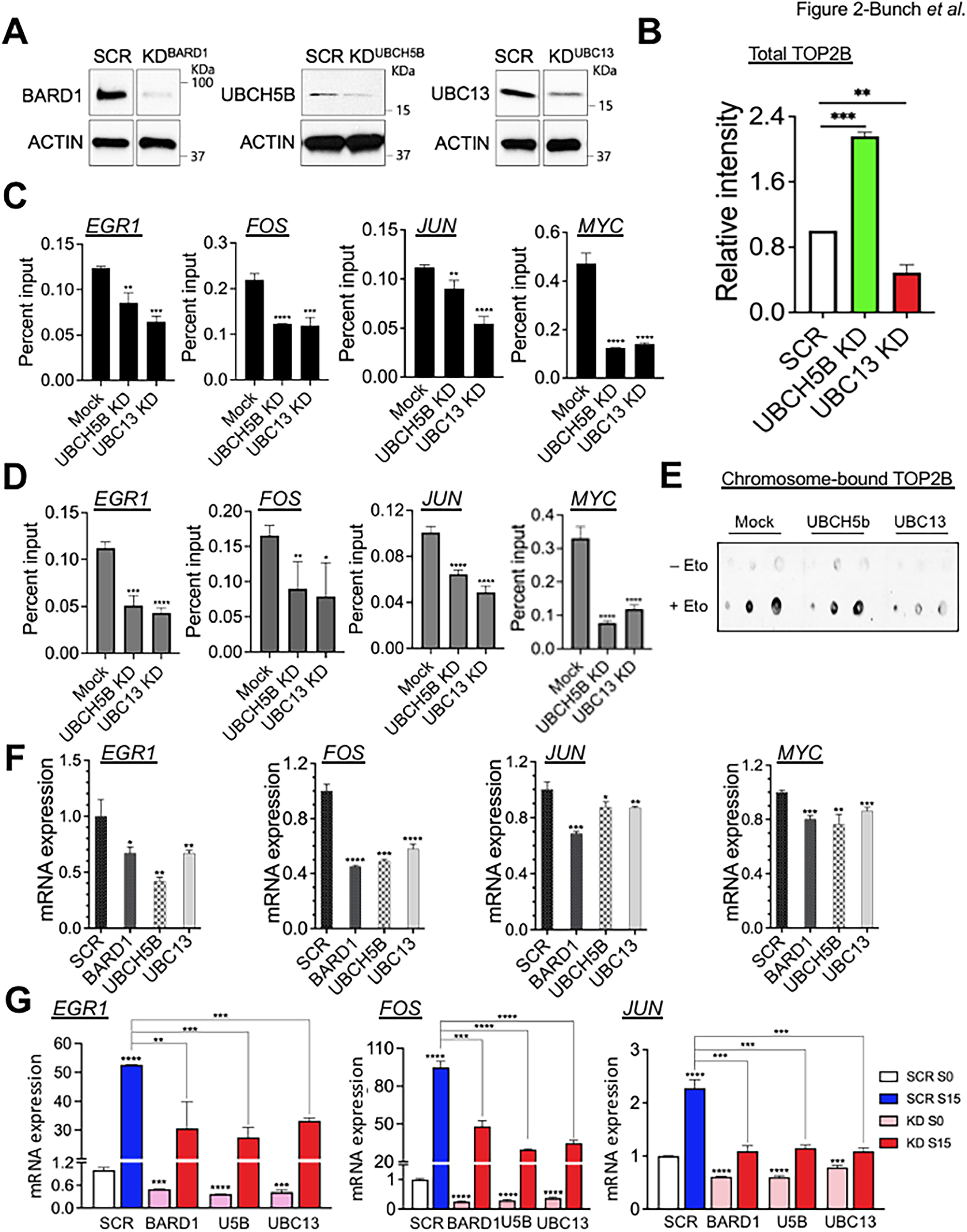
BARD1, UBCH5b, or UBC13 knockdown deregulates TOP2B and represses IEG transcription. (**A**) Immunoblotting results. Left, BARD1 KD validation. Middle, UBCH5b KD validation. Right, UBC13 KD validation. SCR, scrambled, non-specific siRNA control. ACTIN was used as a reference gene and a loading control. (**B**) Summary of multiple immunoblotting results and statistical validations (n = 3 independent experiments). Data are presented as mean values and standard deviations (SDs). *P*-values for the bar graphs were calculated with the unpaired, one sided Student’s t-test. \*\**P* < 0.01, \*\*\**P* < 0.005. (**C**) ChIP-qPCR results showing TOP2B occupancy changes in the TSSs of representative IEGs, *EGR1*, *FOS*, *JUN*, and *MYC*, comparing mock-treated (Mock), UBCH5b KD, or UBC13 KD HEK293 cells (n = 3 independent experiments). Data are presented as mean values and SD. *P*-values for the bar graphs were calculated with the unpaired, one sided Student’s t-test. \*\**P* < 0.01, \*\*\**P* < 0.005, \*\*\*\**P* < 0.0005. (**D**) ChIP-qPCR results showing TOP2A occupancy changes in the TSSs of representative IEGs, *EGR1*, *FOS*, *JUN*, and *MYC*, comparing mock-treated (Mock), UBCH5b KD, or UBC13 KD HEK293 cells (n = 3 independent experiments). Data are presented as mean values and SD. *P*-values for the bar graphs were calculated with the unpaired, one sided Student’s t-test. \**P* < 0.05, \*\**P* < 0.01, \*\*\**P* < 0.005, \*\*\*\**P* < 0.0005. (**E**) Representative ICE assay results showing the effects of UBCH5b or UBC13/MMS2 KD in comparison with the control (Mock). When indicated (+ Eto), etoposide was supplemented to a final concentration of 50 μM for 1 h. The control samples (- Eto) were treated with the same amount of DMSO. Chromosome-bound TOP2B was quantified using immunoblotting. (**F**) qRT-PCR results showing the repressive effects of BARD1, UBCH5b or UBC13 KD at the *EGR1*, *FOS*, *JUN*, and *MYC* genes, compared to the SCR control in HEK293 cells (n = 3 biologically independent samples). Data are presented as mean values and SD. *P*-values for the bar graphs were calculated with the unpaired, one sided Student’s t-test. \**P* < 0.05, \*\**P* < 0.01, \*\*\**P* < 0.005, \*\*\*\**P* < 0.0005. (**G**) qRT-PCR results indicating transcriptional repression in both S0 and S15 conditions at the *EGR1*, *FOS*, and *JUN* genes in HEK293 cells lacking BARD1, UBCH5b, or UBC13 (n = 3 biologically independent samples). Data are presented as mean values and SD. *P*-values for the bar graphs were calculated with the unpaired, one sided Student’s t-test. \*\**P* < 0.01, \*\*\**P* < 0.005, \*\*\*\**P* < 0.0005.

On the other hand, either UBCH5b or UBC13 KD simultaneously decreased TOP2B occupancies at the TSSs of representative IEGs, *EGR1*, *FOS*, *JUN*, and *MYC*, monitored by chromatin-immunoprecipitation-quantitative PCR (ChIP-qPCR)(**Fig. 2C** and **Supplementary Figure 2B**).

Additionally, we monitored TOP2A occupancies at the IEGs in the control (Mock) and KD cells because TOP2A has emerging roles in transcriptional regulation and may play mutual and distinctive roles from ones of TOP2B[1, 67–69]. ChIP-qPCR results, comparing the control and KD cells, showed that TOP2A association with these genes was also compromised, similarly to the case of TOP2B (**Fig. 2D** and **Supplementary Figures 2B–D**), suggesting that BARD1, UBCH5b, and UBC13 may also ubiquitinate and regulate TOP2A. Despite these interesting findings with TOP2A, due to their accumulating redundant and unique, rather complex roles in transcription, we decided to carry out the experiments solely focusing on TOP2B as our main protein of interest in this study.

Genome-bound TOP2B levels were evaluated using the *in vivo* complex of enzymes (ICE) assay. Although UBCH5b KD significantly increased cellular TOP2B protein levels, it did not increase but decreased chromosome-bound TOP2B (**Figs. 2B,E** and **Supplementary Figure 2E**). Consistently with the decreased cellular TOP2B protein levels, UBC13 KD reduced TOP2B-chromosome association (**Figs. 2B,E** and **Supplementary Figure 2E**). We used etoposide, a small chemical inhibitor that traps TOP2 within the normally transient TOP2-DNA cleavage complex[14], and compared genome-wide TOP2B catalytic activity in the control with UBCH5b or UBC13 KD cells. UBC13 KD markedly reduced TOP2B catalytic activities, while only a slight reduction was observed in UBCH5b KD cells (**Fig. 2E**). These results suggested that UBCH5b KD may enhance TOP2B association with the chromosome without critically affecting TOP2B catalysis, whereas UBC13 KD may decrease TOP2B protein stability and catalysis on the chromosomes.

We assessed the effects of BARD1, UBCH5b, or UBC13 KD on IEG transcription by quantitative real-time PCR (qRT-PCR). The data showed that these factors are required for full transcription efficiency at representative IEGs (**Fig. 2F**). The mRNA expression of representative IEGs, *EGR1*, *FOS*, and *JUN*, was compared between the SCR control and KD cells, in serum-starvation (S0) and serum-induction for 15 min (S15). The data indicated that BARD1, UBCH5b, or UBC13 KD interferes with transcription at these genes in both S0 and S15 conditions (**Fig. 2G** and **Supplementary Figures 2B,C**). Overall, our data shown in **Figs. 1** and **2** suggest the distinct patterns and roles of BRCA1-BARD1-mediated TOP2B ubiquitination, determined by E2 enzymes UBCH5b and UBC13/MMS2.

Although both E2 enzymes and BARD1 are important for TOP2B function and IEG transcription, UBCH5b-mediated TOP2B ubiquitination appears to regulate TOP2B-gene association and TOP2B protein degradation, whereas UBC13/MMS2-mediated TOP2B ubiquitination is likely to control TOP2B stability and catalytic activity.

### A transcription-regulatory circuit between BARD1 and mitogen-activated protein kinase pathway

We sought to identify the regulatory mechanisms of the BRCA1-BARD1 complex-mediated TOP2B ubiquitination. To screen the genes, whose expression is directly or indirectly regulated by BARD1, BARD1 KO was generated using CRISPR-Cas9 with gRNAs targeting BARD1 exon 4 (**Supplementary Data 4,5**). As previously reported, a homozygous BARD1 KO was also lethal[56] in our trial, as we could obtain only a heterozygous BARD1 KO (BARD1^−/+^) HEK293 cell line for this work. Generation of the BARD1^−/+^ cell line was confirmed by PCR, genomic sequencing (**Supplementary Data 4,5**), and immunoblotting (**Fig. 3A** and **Supplementary Figure 3A**). It is note that BARD1^−/+^ HEK293 cells seem freeze-thaw sensitive to arrest cell growth after a couple of free-thaw cycles. As described previously, the TOP2B protein level was markedly reduced along with a decreased BARD1 protein level in BARD1^−/+^ cells (**Fig. 3A**). The TOP2A level was also mildly reduced, whereas the γH2AX level was notably elevated in BARD1^−/+^ cells (**Fig. 3A** and **Supplementary Figure 3A**), suggesting accumulated genomic instability in BARD1-deficient cells.

**Fig. 3.**
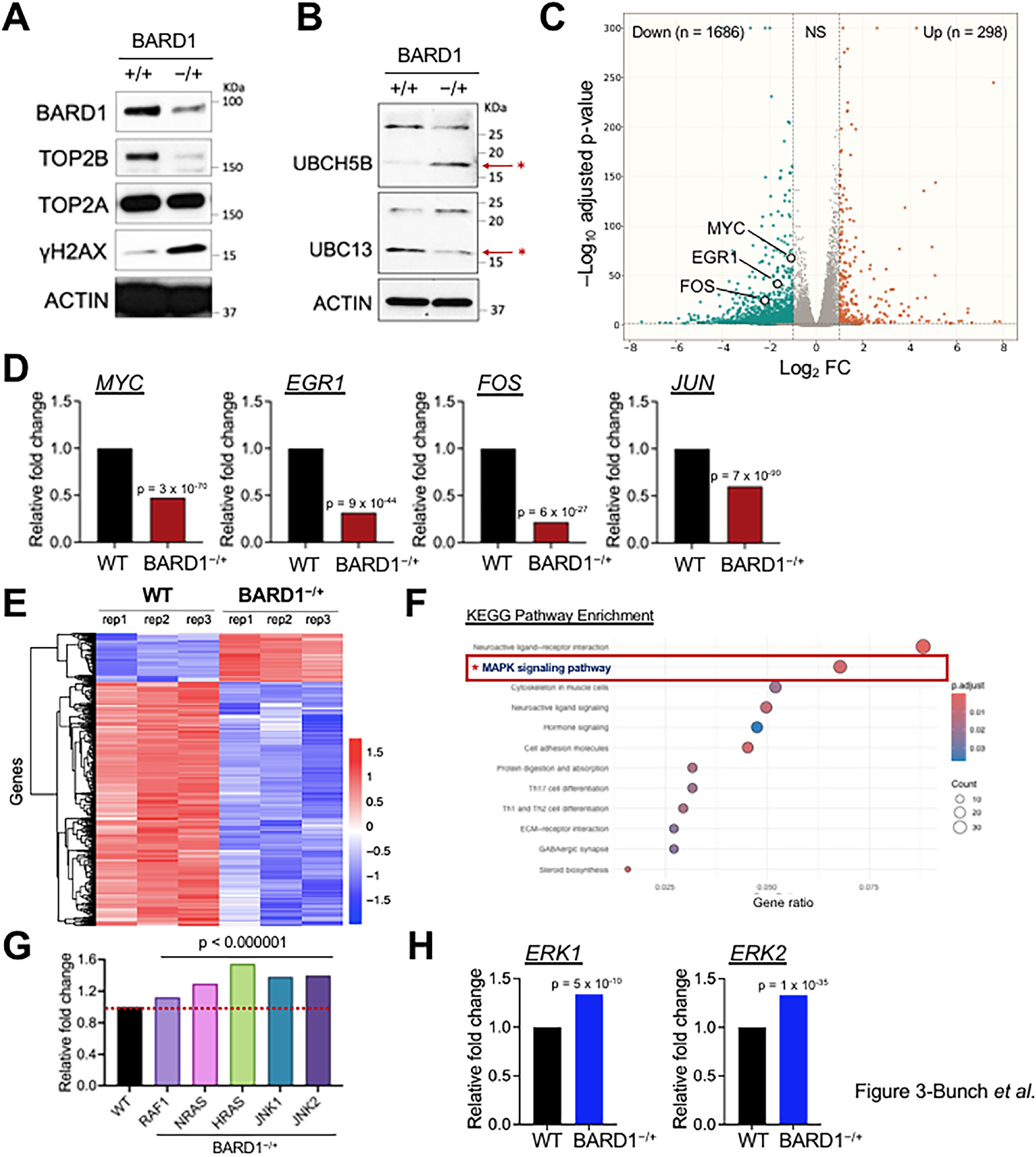
A transcription-regulatory circuit between BARD1 and mitogen-activated protein kinase pathway. (**A**) Immunoblotting results showing protein expression levels of BARD1, TOP2B, TOP2A, and γH2AX in WT (+/+) and BARD1^−/+^ HEK293 cells. ACTIN, a reference gene and a loading control. (**B**) Immunoblotting data showing UBCH5b and UBC13 protein levels in WT and BARD1^−/+^ HEK293 cells. ACTIN, a reference gene and a loading control. (**C**) A volcano plot showing DEGs (|Log_2_FC| > 1 and adjusted *P*-value < 0.05) from RNA-seq data (n = 3), comparing WT and BARD1^−/+^ HEK293 cells. NS, non-significant. (**D**) Relative mRNA expression fold changes of representative IEGs, *EGR1*, *FOS*, *JUN*, and *MYC* in WT and BARD1^−/+^ HEK293 cells. (**E**) Heatmaps of DEGs, comparing WT and BARD1^−/+^ HEK293 cells. (**F**) KEGG pathway analysis with DEGs, comparing WT and BARD1^−/+^ HEK293 cells. (**G**) mRNA expression of key factors, *RAF1*, *NRAS*, *HRAS*, *JNK1*, and *JNK2* in MAPK pathway in WT and BARD1^−/+^ HEK293 cells. (**H**) mRNA expression of *ERK1* and *ERK2* in WT and BARD1^−/+^ HEK293 cells.

Interestingly, we found the opposing effects of BARD1 deficiency on cellular UBCH5b and UBC13 protein levels. In BARD1^−/+^ cells, UBCH5b was increased, while the UBC13 level was diminished (molecular weight of UBCH5b and UBC13, approximately 17 KDa; **Fig. 3B**).

Total RNA-seq comparing WT and BARD1^−/+^ cells revealed that a larger number of genes were downregulated [Log_2_ fold change (FC) < 1; p < 0.05; n = 1686] and a smaller number of genes were upregulated in BARD1^−/+^ cells [Log_2_ fold change (FC) > 1; p < 0.05; n = 298](**Fig. 3C** and **Supplementary Data 6**). We observed the mRNA levels of representative IEGs, including *EGR1*, *MYC*, *FOS*, and *JUN*, were significantly decreased in BARD1^−/+^ cells, consistently with the qRT-PCR results comparing WT and BARD1 KD cells (**Figs. 2F,G**, **3C,D** and **Supplementary Figure 3B**). A heatmap depicting differentially expressed genes (DEGs) upon BARD1^−/+^ was shown in **Fig. 3E**. KEGG pathway analysis projected neuroactive ligand-receptor interaction and the MAPK pathway as the most affected pathways in BARD1^−/+^ cells (**Fig. 3F**). For the neuroactive ligand-receptor interaction, we note that recent studies have indicated an important, bidirectional kidney-brain axis to regulate kidney and brain functions and expression of a few critical neuronal genes in kidney[70, 71]. It was particularly interesting to learn that BARD1 KO affects the MAPK pathway because it is a pathway that is well-known to stimulate the IEG expression. The expression of critical MAPK pathway genes[40], including *RAF1*, *NRAS*, *HRAS*, *JNK1*, *JNK2*, *ERK1*, and *ERK2*, was monitored, comparing WT and BARD1^−/+^ cells. Intriguingly, these key constituents of the MAPK pathway were mildly yet significantly upregulated in BARD1^−/+^ cells (**Figs. 3G,H** and **Supplementary Data 7**). It seemed paradoxical that BARD1 deletion could cause the increased expression of key genes in MAPK pathway, while decreasing critical target genes of the MAPK pathway (**Figs. 3C,D,G,H** and **Supplementary Figure 3B**). BARD1 is a DNA binding protein (**Supplementary Figure 3C**) and stably associated with IEG promoters[29]. Based on the unexpected findings and spatial and temporal relations between BARD1 and ERKs, we hypothesized whether BARD1 could be a downstream effector regulated by these MAPK pathway proteins, to control the expression of IEGs.

### Activated ERK2 phosphorylates BARD1 at S391, a pathologically important residue for IEG activation

To test this hypothesis, BARD1 protein was expressed and purified from HEK293F cells using the YFP-tag system[58, 59]. Human ERK1 and ERK2 and well-characterized, constitutively active R84S ERK1 (ERK1m) and R67S ERK2 (ERK2m) mutants[14, 72] were purified from bacteria as described in our previous work[14]. We performed an *in vitro* kinase assay using these proteins with ^32^P-radioactive ATP. We found that BARD1 was phosphorylated by ERK1m and ERK2m (**Fig. 4A**). Notably, ERK1 could be auto-phosphorylated, which was a previously unknown phenomenon, while ERK2 could not, and both unmodified ERK1 and ERK2 did not phosphorylate BARD1 (**Fig. 4A**). Next, we performed the *in vitro* kinase assay with cold ATP, and excised the bands corresponding to BARD1 on an SDS PAGE. Control (ERK storage buffer only), BARD1-ERK1m, and BARD1-ERK2m reactions, were subsequently analyzed by phospho-peptide enrichment LC–MS/MS analysis (**Figs. 4B,C**). The mass spectrometry data identified residue S394 in the control BARD1 and a phosphorylation of S391 uniquely by ERK2m with high confidence (**Figs. 4D,E**, **Supplementary Figure 4A**, and **Supplementary Data 8–12**). Interestingly, the residue S391 has been reported as a mitotic BARD1 phosphorylation site and its mutations have been implicated in cancers[56, 73]. The kinase has been unknown so far, and to our best knowledge, this is the first study that unveiled that activated ERK2 is responsible for BARD1 phosphorylation at S391 (**Figs. 4A–E**, **Supplementary Figure 4A**, and **Supplementary Data 8–12**).

**Fig. 4.**
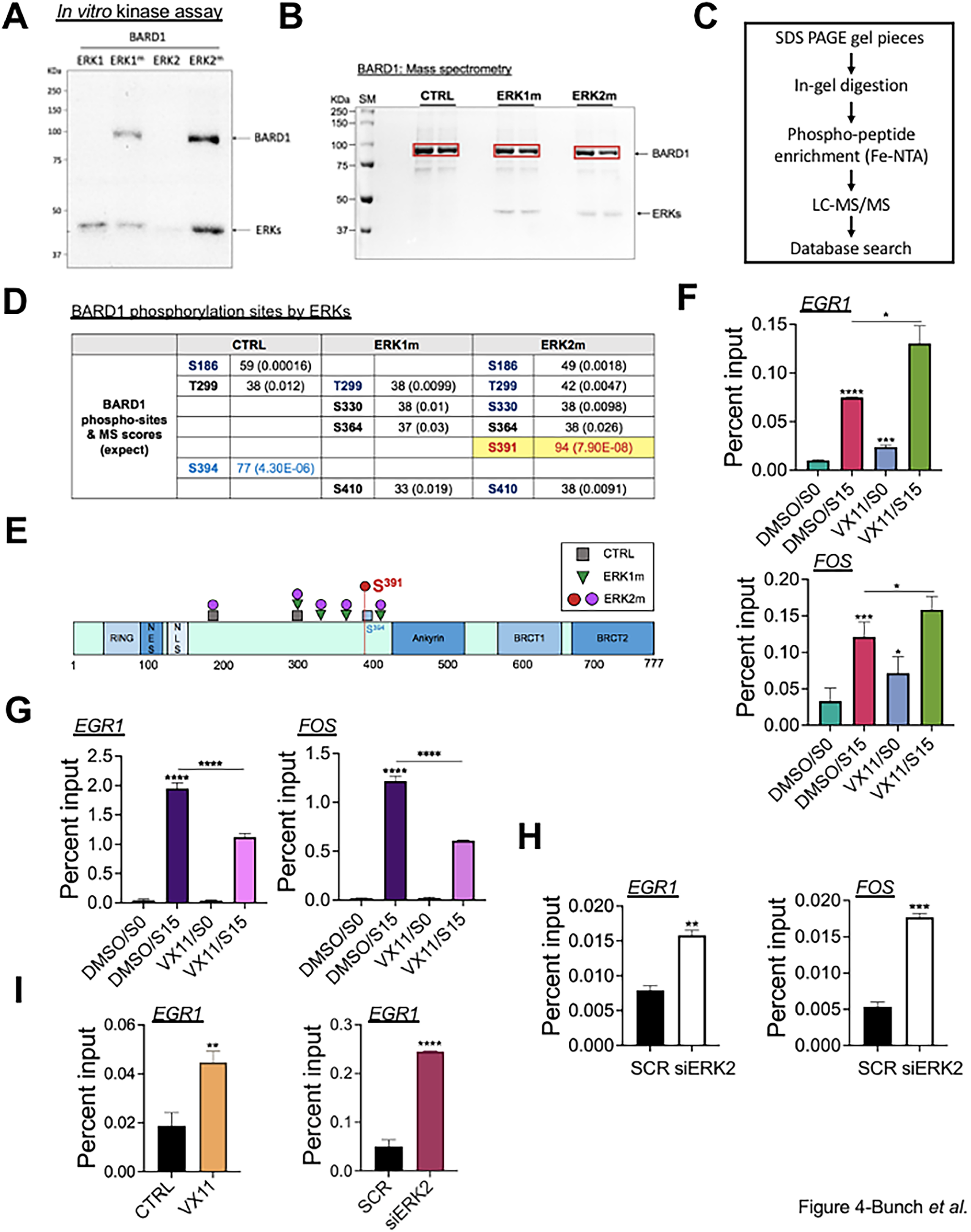
Activated ERK2 phosphorylates BARD1 at S391, a pathologically important residue for IEG activation. (**A**) Autoradiograms of the *in vitro* kinase assay with BARD1 and ERKs. (**B**) *In vitro* kinase assay followed by SDS-PAGE for the preparation of BARD1 samples for mass spectrometry analyses. Red boxes showing sliced gel pieces including BARD1 used for the analyses. CTRL, ERK storage buffer only without ERKs; SM, size marker (KDa). (**C**) A flow chart showing the major steps of mass spectrometry analyses performed in this work. (**D**) Summary of the phosphorylated residues of BARD1 in the control, ERK1m, and ERK2m reactions identified by mass spectrometry analyses and their scores. The residues with high confidence and statistical significance are marked with colors. (**E**) A schematic representation of the BARD1 residues phosphorylated by ERK2m (a red and purple circles) and ERK1m (green triangles) and in the control (a skyblue and gray squares). The residues identified with high confidence were marked with a skyblue square (S394 in the control BARD1) and a red circle (S391 in BARD1 phosphorylated by ERK2m). (**F**) ChIP-qPCR results showing BARD1 occupancy changes in the TSSs of representative IEGs, *EGR1* and *FOS*, comparing DMSO- or VX11-treated cells in S0 and S15 conditions (n = 3 independent experiments). Data are presented as mean values and SD. *P*-values for the bar graphs were calculated with the unpaired, one sided Student’s t-test. \**P* < 0.05, \*\*\**P* < 0.005, \*\*\*\**P* < 0.0005. (**G**) ChIP-qPCR results showing S2 Pol II occupancy changes in the TSSs of representative IEGs, *EGR1* and *FOS*, comparing DMSO- or VX11-treated cells in S0 and S15 conditions (n = 3 independent experiments). Data are presented as mean values and SD. *P*-values for the bar graphs were calculated with the unpaired, one sided Student’s t-test. \*\*\*\**P* < 0.0005. (**H**) ChIP-qPCR results showing BARD1 occupancy changes in the TSSs of representative IEGs, *EGR1* and *FOS*, comparing SCR control and ERK2 KD HEK293 cells in S0 and S15 conditions (n = 3 independent experiments). Data are presented as mean values and SD. *P*-values for the bar graphs were calculated with the unpaired, one sided Student’s t-test. \*\**P* < 0.01, \*\*\**P* < 0.005. (**I**) ChIP-qPCR results showing TOP2B occupancy changes in the TSSs of *EGR1* gene, comparing DMSO- or VX11-treated cells (left) and SCR and ERK2 KD cells (right)(n = 3 independent experiments). Data are presented as mean values and SD. *P*-values for the bar graphs were calculated with the unpaired, one sided Student’s t-test. \*\**P* < 0.01, \*\*\*\**P* < 0.0005.

We investigated the effect of ERK2-mediated BARD1 phosphorylation on IEG transcription and BARD1-gene interaction. VX-11e (VX11), a small chemical inhibitor specific to ERK2[74, 75], was employed, and the occupancies of BARD1 and phosphorylated Pol II at S2 of the RPB1 subunit (S2 Pol II) were monitored in serum-starvation and -induction, S0 and S15 conditions. VX11 was supplemented to the cells at a final concentration of 100 nM for 3 h with the control dimethyl sulfoxide (DMSO) side-by-side. Strikingly, ERK2 catalytic inhibition by VX11 significantly increased the BARD1 level associated with the TSSs of *EGR1* and *FOS* (**Fig. 4F**). By contrast, upon serum-induced gene activation, S2 Pol II levels were notably decreased in the gene bodies of *EGR1* and *FOS* in the presence of VX11 (S15; **Fig. 4G**). Additionally, we depleted ERK2 using siRNA species targeting ERK2 and found that ERK2 KD increased BARD1 occupancies in the TSSs of *EGR1* and *FOS*, similarly to the results from ERK2 catalytic inhibition (**Fig. 4H** and **Supplementary Figure 4B**). As shown in our previous study[14], VX11 treatment or ERK2 KD increased TOP2B occupancies in the TSSs of *EGR1* and *FOS* genes (**Fig. 4I**). Together, these data suggested that ERK2-mediated phosphorylation of BARD1 is required for proper BARD1 association and function and transcriptional activation in IEGs.

### BARD1 phosphorylation by activated ERK2 enhances UBCH5b- and UBC13/MMS2-mediated TOP2B ubiquitination

To define the mechanistic basis of ERK2-mediated BARD1 phosphorylation in TOP2B ubiquitination by BRCA1-BARD1 with UBCH5b or UBC13/MMS, each of the BRCA1-BARD1 complex, TOP2B, and ERK2m proteins was prepared through a series of liquid chromatography procedures, as previously established (**Fig. 5A**)[14, 29] and validated by immunoblotting (**Fig. 5B**). These proteins along with E1 and two E2 enzymes, UBCH5b and UBC13 were utilized for the *in vitro* ubiquitination assay. To inhibit ERK2m kinase activity, VX11 was supplemented to the designated reactions at 1 μM final concentration. E1 (approximately 120 KDa) and BRCA1-BARD1 (220 KDa and 100 KDa) as the E3 ligase were added in minimal amounts (1 nM for BRCA1-BARD1; 50 nM for E1 enzyme) to avoid signal overlapping as their molecular weights are similar to that of TOP2B (180 KDa). Approximately 22% of each terminated reaction was visualized by SDS-PAGE and silver-staining, validating equivalence among reactions and the absence of visible protein degradation (**Fig. 5C**). TOP2B ubiquitination and TOP2B were monitored by immunoblotting. The results revealed enhanced TOP2B ubiquitination by BRCA1-BARD1-UBCH5b in the presence of ERK2m, evidenced by smearing in upper bands including a prominent band in a high molecular weight region (red boxes for comparison and a red asterisk for the prominent band, **Fig. 5D**). BRCA1-BARD1-UBC13/MMS2 robustly ubiquitinated TOP2B, and the presence of ERK2m apparently further enhanced TOP2B ubiquitination by the complex (light blue boxes for comparison, **Fig. 5D**). VX11 inhibited ERK2m-stimulated TOP2B ubiquitination, to the levels similar to the control reactions without ERK2m (yellow boxes, **Fig. 5D**), further confirming the novel function of activated ERK2 in stimulating BRCA1-BARD1 coupled with UBCH5b or UBC13/MMS2 to ubiquitinate TOP2B.

**Fig. 5.**
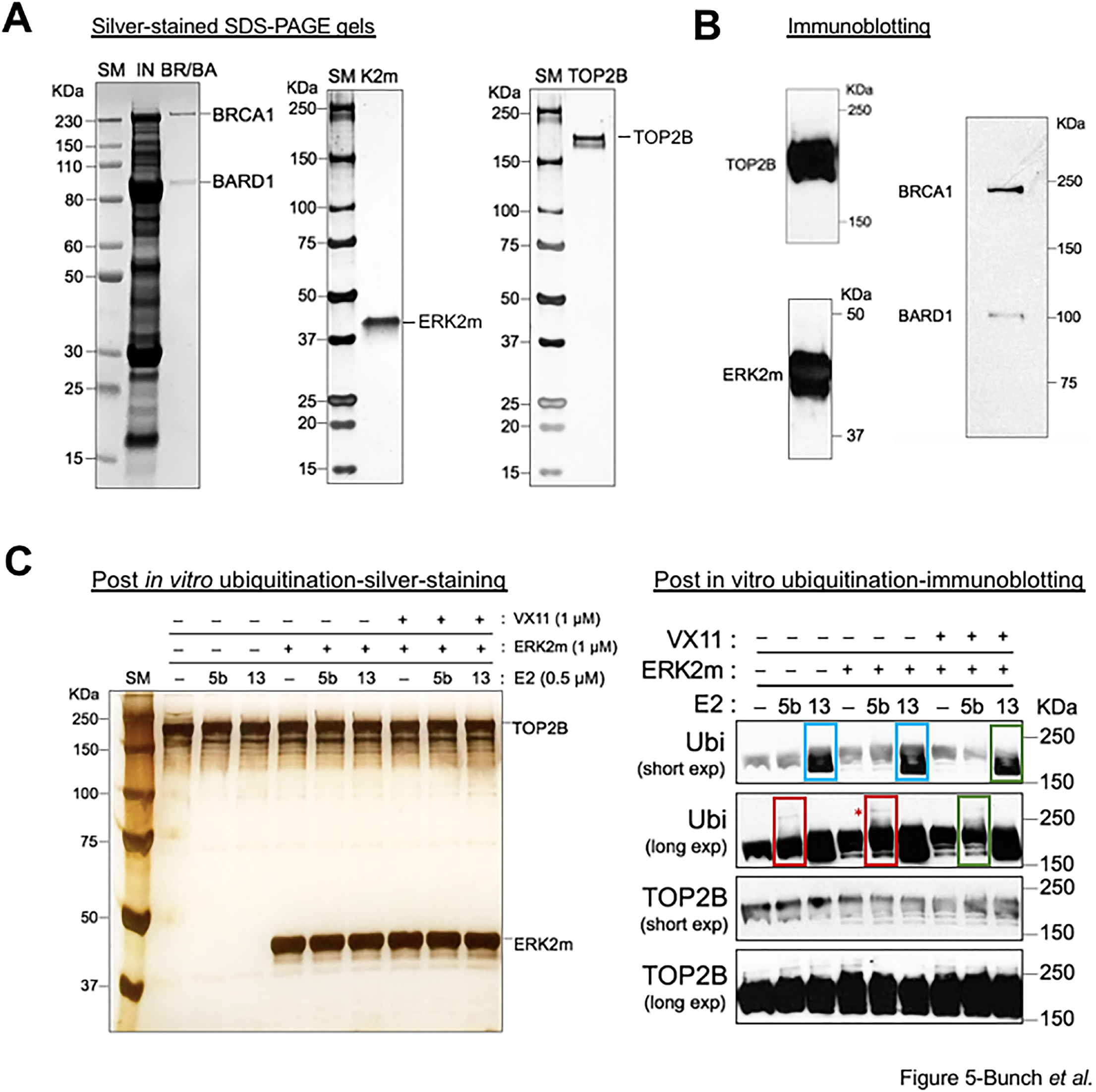
BARD1 phosphorylation by activated ERK2 enhances UBCH5b- and UBC13/MMS2-mediated TOP2B ubiquitination. (**A**) Silver-stained SDS PAGE gels showing BRCA1-BARD1 purified as a complex from HEK293 cells (left), ERK2m purified from E. coli (middle), and TOP2B from HEK293 cells (right). IN, input protein mixtures including BRCA1-BARD1 loaded onto a sizing exclusion column. (**B**) Immunoblotting results verifying TOP2B (upper left), ERK2m (bottom left), and BRCA1-BARD1 (right). (**C**) Silver-stained SDS-PAGE gel showing a portion (22%) of the reactions immediately after the *in vitro* ubiquitination assay. (**D**) Immunoblotting results showing TOP2B ubiquitinated by UBCH5b-BRCA1-BARD1 (red boxes) or UBC13/MMS2-BRCA1-BARD1 (skyblue boxes) in the absence and presence of ERK2m. A band, corresponding presumably to a higher molecular-weighted, poly-ubiquitinated TOP2B, with UBCH5b-BRCA1-BARD1 becomes more intense in the presence of ERK2m (marked with a red asterisk). VX11 supplement along with ERK2m interferes with stimulated ERK2m-mediated TOP2B ubiquitination by UBCH5b-BRCA1-BARD1 and UBC13/MMS2-BRCA1-BARD1 (green boxes). Ubiquitin (Ubi) and TOP2B blots were developed for a longer (long exp) and shorter (short exp) period for better presenting the bands.

## DISCUSSION

Metazoan stress-inducible gene regulation employs a systematic regulatory step downstream of the TSS, immediately before productive elongation, called Pol II pausing and pause release[15, 76]. This step ensures a prompt and synchronized transcriptional burst to maximize the efficiency of gene regulation, for genes that must respond rapidly to cellular signals. We have been studying the mechanisms of Pol II pausing and pause release, using representative Pol II pausing model genes, such as *HSP70*, hypoxia-inducible genes (HIGs), and IEGs[10, 11, 14, 29, 30, 77]. In particular, *HSP70*, *EGR1*, and *FOS* display well-established, prominent Pol II pausing and their expression is profoundly dependent on complex regulatory networks that convert paused Pol II into a processive enzyme. For example, the catalytic activity of TOP2B and DDR signaling are required for productive transcription in IEGs[1, 14, 30, 41]. Inhibiting TOP2B or DNA-PK using small chemical inhibitors prevents Pol II from being released from pausing and hinders transcriptional activation in IEGs.

Whether TOP2B causes DSBs as a programmed, gene regulatory custom or as a random, accidental event remains as debatable, however, it has been shown that TOP2B-mediated DSBs activate transcription in diverse sets of genes[14, 30, 43, 78, 79]. For example, etoposide, trapping TOP2B after the generation of DSB, stimulates IEG transcription, whereas ICRF193, a TOP2B catalytic inhibitor, represses it[14, 30]. The proposed sequence of events is that TOP2B is modulated by a transcription factor(s) or DNA topological stresses that activate TOP2B catalytic activity; TOP2B mediates DSB to condition DNA topology that is favorable for transcription; DSB provokes DDR signaling; DNA repair factors are activated, which is an important question for future studies to clarify.

To understand the upstream factors or events that regulate TOP2B binding to genes and catalysis, we have been investigating key transcription factors implicated in Pol II pause release, including BRCA1-BARD1, DNA-PK, and ERKs[1, 11, 14, 29, 30]. Among these factors, BARD1, TOP2B, and DNA-PK are abundantly associated with TSSs of the genes harboring Pol II pausing during the transcriptionally resting state[29, 30]. Previously, it was shown that BRCA1-BARD1 ubiquitinates TOP2B and contributes to increased TOP2B stability and affinity for DNA in the Pol II pausing state[29]. Transcriptional activation is coupled with DDR signaling in IEGs, leading to the activation of ATM and phosphorylation of BRCA1 at S1524, γH2AX, and DNA-PK[1, 29, 30, 41]. Furthermore, ERK2 can directly phosphorylate TOP2B to control TOP2B catalysis of positive and negative DNA supercoiling[14]. Interestingly, ERK2 promotes TOP2B to resolve positive supercoiling and yet, delays it to relax negative DNA supercoiling[14].

The findings of this work deepen the mechanistic understanding of the dynamic interactions and regulations among these important transcriptional, DNA repair, and DNA topological factors (**Fig. 6**). First, we found that BRCA1-BARD1-mediated TOP2B ubiquitination has differential effects on TOP2B activity depending on the engaging E2 enzymes, UBCH5b vs UBC13/MMS2 (**Figs. 1A–E**, **2B,E,5D** and **Supplementary Figures 1A–D,2A**). UBCH5b is a ubiquitously expressed E2 known to mediate K48-linked polyubiquitination, and promotes protein destabilization and DNA repair [80],[81, 82], whereas UBC13/MMS2 is a more specialized E2 that synthesizes K63-linked polyubiquitination for signal transduction and DNA repair[83, 84]. UBCH5b KD elevated cellular TOP2B protein levels without affecting its genome activities (**Figs. 2B,E** and **Supplementary Figure 2A**), while compromising IEG transcription (**Figs. 2F,G**), which suggests that UBCH5b-mediated TOP2B dissociation/destabilization may be important for proper IEG transcriptional activation. UBC13 KD decreased both cellular protein levels and the genome-wide activity of TOP2B as well as IEG transcription (**Figs. 2B,E,F,G** and **Supplementary Figures 2A**), suggesting that UBC13-mediated TOP2B ubiquitination could be required for TOP2B stability and catalysis in IEG transcription.

**Fig. 6.**
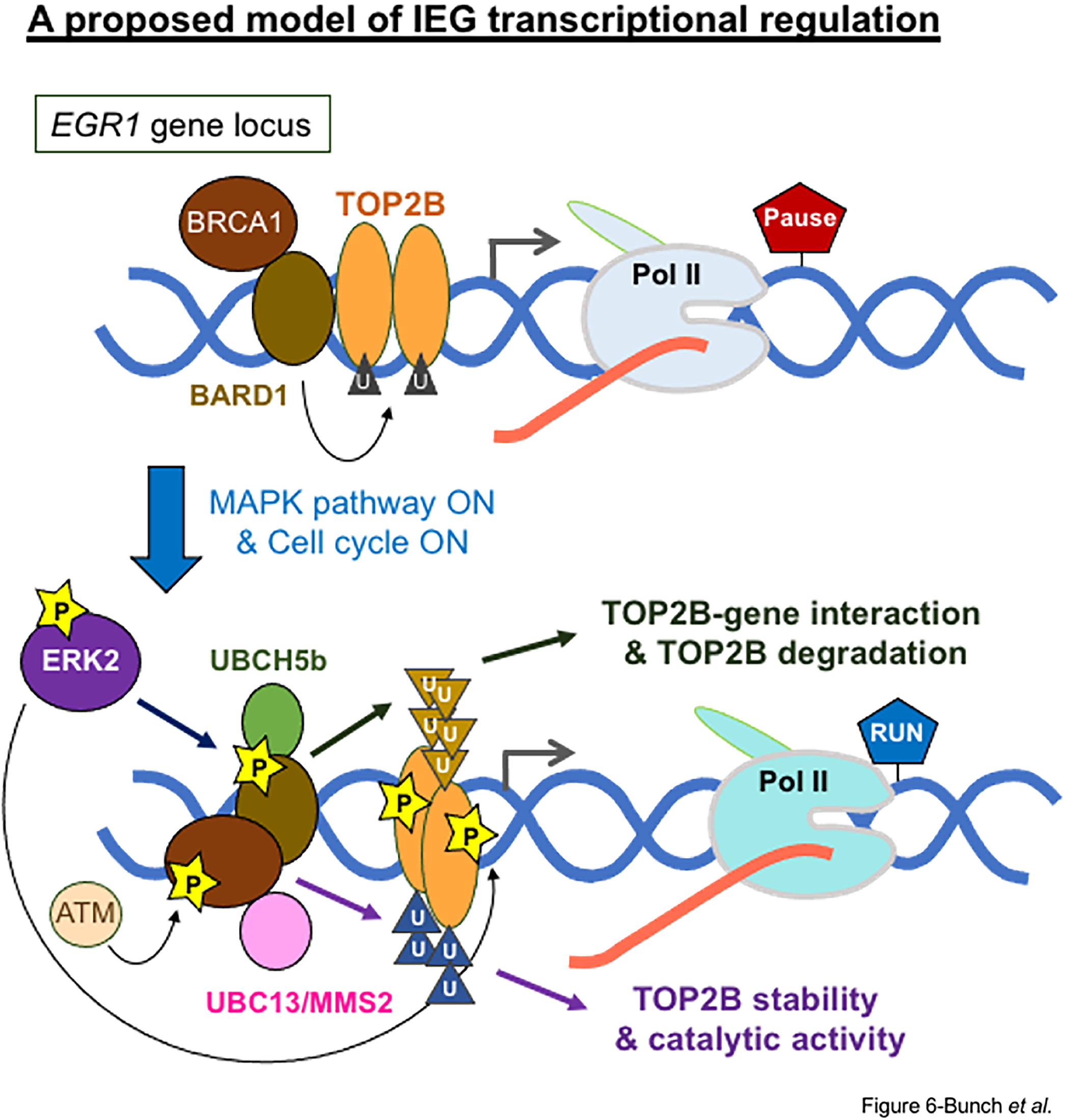
A proposed model: the triangular relationship of TOP2B, BARD1, and ERK2 for human IEG transcription. Based on the findings, we propose a gene regulatory model involving dynamic interactions among a critical DNA topology regulator, a DNA repair factor, and a transcriptional activator, TOP2B, BRCA1-BARD1, and ERK2m in IEG transcription. In Pol II pausing during the transcriptionally resting state, BRCA1-BARD1 ubiquitinates TOP2B, which increases TOP2B stability and affinity with the IEG TSSs[1, 29]. Upon the reception of a signal to progress the cell cycle to the G1 phase, ERK2 is phosphorylated and activated in the cytoplasm and translocates into the nucleus[40]. Activated ERK2 phosphorylates BARD1 at S391 (**Figs. 4A–E**) and other previously reported substrates, TOP2B, and ELK1[14]. BARD1/TOP2B phosphorylation by ERK2 and TOP2B ubiquitination by ERK2-stimulated UBCH5b-BRCA1-BARD1 and UBC13-BRCA1-BARD1 are important for proper BARD1/TOP2B engagement and productive IEG transcription (**Figs. 2C,E–G, 3C, D, 4A–I, 5D**). Our data suggest that TOP2B ubiquitination by UBCH5b-BRCA1-BARD1 may promote TOP2B-gene interaction and TOP2B degradation whereas TOP2B ubiquitination by UBC13/MMS2-BRCA1-BARD1 may enhance TOP2B catalysis and stabilize TOP2B (**Figs. 2A–C,E**). Pol II pause release is coupled with TOP2B catalysis and DDR signaling, which involves the activation of DNA repair factors including ATM, ATR, and DNA-PK[1, 30, 41].

Second, our work identified a bidirectional relationship between the MAPK pathway and BARD1 and discovered activated ERK2 as a novel kinase that phosphorylates BARD1 at S391 (**Figs. 3E–H**, **4A–E**). Recent studies have emphasized the functional crosstalk between the DDR and MAPK pathways for the determination of cell fate and its abnormality in cancers[85, 86]. It is notable that BARD1 KD markedly increases cellular γH2AX levels (**Fig. 3A** and **Supplementary Figure 3A**), implying the accumulation of DNA damage that can contribute to the activation of key players in the MAPK pathway. Additionally, our data suggests an important role of BARD1 and BARD1-mediated ubiquitination of its substrates (e.g., TOP2B) for IEG transcription in response to a MAPK pathway activation, because IEG expression is diminished without BARD1, despite significantly increased levels of key MAPK pathway genes (**Figs. 2A,F,G, 3A,C,D** and **Supplementary Figure 3B**). Third, activated ERK2-mediated BARD1 phosphorylation stimulates TOP2B ubiquitination by both E2–E3 complexes, UBCH5b-BRCA1-BARD1 and UBC13/MMS2-BRCA1-BARD1 (**Fig. 5D**). Inhibiting ERK2 results in abnormal increases/retentions of BARD1 and TOP2B and transcriptional repression in IEGs (**Figs. 4F–I**). We suggest that the stimulation of BARD1-mediated TOP2B ubiquitination by activated ERK2, in conjunction with UBCH5b and UBC13/MMS2, is important for proper TOP2B dissociation/mobilization and catalysis and thus for transcriptional activation.

Recent studies by others and our own have shown distinctive Pol II pausing regulatory mechanisms in HIGs in comparison to what has been understood in IEGs[1, 11, 87]. The most critical difference is that the DDR signaling seems to rarely be involved in Pol II pause release in HIGs, although similar DNA repair factors play key roles in Pol II pausing and pause release, including DNA-PK, TRIM28, and TOP2B in both IEGs and HIGs[30, 87]. These gene-specific differential mechanisms may be explained by genomic context-dependent DNA topological signatures and requirements[1, 11]. For instance, TOP2B inhibition by small chemicals has different effects on IEGs and HIGs[1, 11, 14, 30]. In HIGs, TOP2B is phosphorylated at T1403 by DNA-PK in the state associated with paused Pol II, where the activity of TOP2B plays an important role in suppressing HIG transcription in the resting state, under normoxia[11]. In contrast, in IEGs, TOP2B catalysis is activated upon transcriptional activation[14, 29, 30, 43]. For example, activated ERK2 phosphorylates and catalytically activates BARD1 and TOP2B, as shown previously and in this study (**Figs. 4F–I**, **5D** and **Supplementary Figure 4A**)[14]. The exact role of DNA-PK phosphorylation/activation in transcriptional activation of subsets of genes such as HIGs, IEGs, *HSP70*, and androgen- and estrogen-receptor genes remains to be uncovered[10, 11, 30, 78, 79, 87, 88]. In particular, BRCA1-BARD1 and DNA-PK reportedly compete for distinctive DSB repair pathways to be engaged with the DSB sites[89, 90]. Is this the reason why these factors co-occupy IEGs, or are there other unknown molecular dynamics involving them? Moreover, it will be interesting to understand what mediates and regulates specific E2 enzymes to interact with BRCA1-BARD1 during Pol II pausing and pause release in IEG transcription. These are some of the critical questions awaiting future investigations.

## CONCLUSIONS

In summary, this study revealed the important role of E2 enzymes in human IEG expression. UBCH5b and UBC13/MMS cooperate with the BRCA1-BARD1 E3 ligase to ubiquitinate TOP2B and yet, in distinct manners and for distinct regulations/functions. Our data suggest that BRCA1-BARD1-UBCH5b regulates TOP2B-gene association/dissociation and TOP2B degradation, while BRCA-BARD1-UBC13/MMS2 controls TOP2B catalytic activity and stability, both of which are important for productive Pol II transcription in IEGs. This study revealed that activated ERK2 is the kinase responsible for BARD1 phosphorylation at S391, reportedly a mitotic phosphorylation and pathologically important site. Activated ERK2-mediated BARD1 phosphorylation stimulates BRCA1-BARD1, in conjunction with UBCH5b and UBC13/MMS2, to ubiquitinate TOP2B, which is required for proper BARD1 and TOP2B behavior and engagement with IEGs, thus for effective transcription and Pol II pause release in human IEGs.

## List of abbreviations

IEG: Immediate Early Gene
TOP2B: Topoisomerase IIβ
mRNA: Messenger RNA
ncRNA: Non-coding RNA
Pol II: RNA Polymerase II
PTM: Post-Translational Modifications
DSB: Double-Stranded DNA Breakage
DDR: DNA Damage Response
ATM: Ataxia Telangiectasia Mutated
ATR: Ataxia Telangiectasia and Rad3-Related
DNA-PK: DNA-dependent Protein Kinase
KD: KnockDown
KO: KnockOut
LC-MS/MS: Liquid Chromatography-Mass Spectrometry/Mass Spectrometry
TSS: Transcription Start Site
MAPK: Mitogen-Activated Protein Kinase
NTD: N-Terminal Domain
CTD: C-Terminal Domain
GFP: Green Fluorescent Protein
SD: Standard Deviation
ChIP-qPCR: Chromatin-ImmunoPrecipitation-quantitative PCR
siRNA: Small Interfering RNA
SCR: Scrambled siRNA
ICE: In vivo Complex of enzymes
qRT-PCR: Quantitative Real-Time PCR
DMSO: DiMethyl SulfOxide
HIG: Hypoxia-Inducible Gene

## Acknowledgments

We are grateful to D. Engelberg and N. Soudah at the Hebrew University of Jerusalem for their generosity in sharing the bacterial WT and mutant human ERK1/2 expression vectors, to J. Chang’s laboratory at Kyungpook National University (KNU) for purifying the ERK proteins, to Rokit Genomics for performing RNA-sequencing analysis, to Enago for proofreading the manuscript, to R. Nakagawa at RIKEN for performing LC–MS/MS analyses, and to Ubigene Biosciences for generating and validating the BARD1 heterozygous KO HEK293 cells for this work. We thank H. Cho at Ajou University Medical School for sharing critical resources and materials. We also thank D. Kim, C. Jung, S. Ju, and the current members of the Bunch laboratory members at KNU for their technical assistance. H.B. thanks R. S. Baker, John and D. Bunch, and J. Christ for their loving encouragement and support throughout the course of this work. This research was supported by grants from the National Research Foundation (NRF) of the Republic of Korea (2022R1A21003569, RS-2025-02303149, and RS-2025-16067324) to H.B.

## Author Contributions

HB performed cell culture, drug treatment, siRNA-mediated factor KD, ChIP-qPCR, total RNA preparation for sequencing, ICE assay, and immunoblotting. HB and JhJ performed qRT-PCR, *in vitro* kinase and ubiquitination assays and prepared for mass spectrometry samples. AC and MS purified and prepared for BARD1, BRCA1-BARD1, and TOP2B proteins. JyJ, IJ, KK performed bioinformatics analysis of RNA-seq data. HB created the hypothesis, designed and coordinated the experiments, analyzed and curated the data, and wrote and revised the manuscript.

